# Protein Plasticity and its Role in Cellular Functions

**DOI:** 10.1101/2020.08.18.256230

**Authors:** Sidra Ilyas, Abdul Manan

**Affiliations:** Dept. of Microbiology and Molecular Genetics, University of the Punjab, Quaid-e-Azam Campus, Lahore 54590, Pakistan; Institute of Molecular Biology and Biotechnology (IMBB), The University of Lahore, Pakistan

**Keywords:** Cysteines, redox, intrinsically disorder region, plasticity, fuzzy complexes, conformational heterogeneity

## Abstract

The contribution of redox active properties of cysteines in intrinsically disordered regions (IDRs) of proteins is not very well acknowledged. Despite of providing structural stability and rigidity, intrinsically disordered cysteines are exceptional redox sensors and the redox status of the protein defines its structure. Experimental evidence suggests that the conformational heterogeneity of cysteines in intrinsically disordered proteins (IDPs) is related to numerous functions including regulation, structural changes and fuzzy complex formation. The unusual plasticity of IDPs make them suitable candidate to interact with many clients under specific conditions. Binding capabilities, dimerization and folding or unfolding nature of IDPs upon interaction with multiple clients assign distinct conformational changes associated with disulfide formation. Here we are going to focus on redox activity of IDPs, their dramatic roles that are not only restricted to cellular redox homeostasis and signaling pathways but also provide antioxidant, anti-apoptotic, binding and interactive power.

## 1. Introduction

The discovery of proteins that function without a well-defined tertiary structure has challenged the sequence-structure-function paradigm. As the tertiary structure of protein is usually achieved by folding it has been investigated that a large proportion of proteins do not mediate folding due to lack of sufficient hydrophobic amino acids and function in a disordered state [1,2]. Therefore, these proteins make up (∼40%) of eukaryotic proteome are known as intrinsically disordered proteins (IDPs) and function well without achieving a defined tertiary structure [3–6] and [7–9]. *In vitro*, IDPs are enriched in regions with greater number of polar disorder-promoting amino acids and depleted in order-promoting amino acids hence, showing greater conformational plasticity [10]. The net charge distribution pattern and low mean hydrophobicity characterize their fully extended conformation (alternative positively and negatively charged residues) or compact globule partially folded conformation (stretches of positively and negatively charged residues) or somewhere in between [11–13].

Structural flexibility of IDPs play numerous roles in regulating cell processes such as signaling, transcription regulation, aggregate formation, DNA condensation, mRNA processing and apoptosis [14]. Under oxidative conditions, at least two cysteines are desired to form a disulfide bridge to fold, providing protein to function properly. Protein containing cysteines as a norm are considered to be in the ordered region. Nonetheless, cysteines residues in the disordered regions of proteins are exceptional and their diverse roles, functions and conformational heterogeneity due to plasticity are not very well investigated. Some of their common features of IDPs are as follows.

## 2. Protein folding and unfolding

In response to specific environmental changes such as stress, ligand(s) and upon binding to different interaction partners, IDPs undergo order-to-disorder and/or disorder-to-order conformational change [15–17]. They have the tendency to undergo binding-induced folding or remain significantly disordered upon binding with another protein resulting in heterogeneous “fuzzy” complex formation [18,19]. IDPs have enormous potential to interact with and control multiple binding clients simultaneously by adopting different conformations interestingly, this structural plasticity and adaptability seems to be an evolutionary conserved feature as numerous research studies reveal that any alteration/mutation in the intrinsically disordered regions (IDRs) would lead to neurodegenerative diseases and cancer [20–22].

## 3. Flexible binding segments enhance plasticity

Various protein clients fuse with IDPs motifs providing conformational heterogeneity in a functionally promiscuous manner. The resulting proteins may have ordered and disordered regions in variable stochiometric ratios which synergistically increase their functional versatility. Conformational plasticity of flexible binding segments on IDPs acts as recognition molecules for DNA (leucine zipper protein-GCN4), RNA (ribosomal proteins e.g., L11-C76) and protein[23,24]. IDPs fusion generated proteins can homo-, hetero-dimerize and/or multimerize via self- and multivalent interactions, become activated and processed due to newly generated sites for modification. Their activation is modulated by complex post-translational modifications (PTM) that involve differential splicing or phosphorylation at specific sites and/or recruitment into hybrid activation complexes. Motifs or PTM sites embedded within IDRs can affect critical cellular processes, increased more complexity and phenotypes during evolution as compared to ordered structures.

## 4. Signaling and regulation

IDPs are engaged in signaling of proteins, kinases, and regulatory networks in the form of transcription factors and splicing factors [25,26]. Diverse PTM of residues within the IDR facilitates the regulation of protein function by encoding and decoding information[27,28]. These unique properties support accomplishing regulatory and signaling functions. Splicing of disordered segments without affecting structured domain(s) can have important consequences in remodeling of signaling and regulatory networks during development. This plasticity of alternative splicing may lead to the emergence of novel proteins between different organisms.

## 5. Aggregate formation

IDPs have a generic tendency to transform their soluble states into insoluble highly structured aggregates or fibrils. Aggregation in such scenarios creates microscopic organization inside the cell under both physiological and pathological conditions by forming higher-order structures having special modifications and interactions. The reversible/irreversible aggregates might interfere with signaling pathways under pathological conditions e.g. fibril formation in prion protein at N terminus [29].

## 6. Intrinsically disordered proteins (IDPs) as a redox sensor

Hybrid protein complexes formed when intrinsically disordered cysteine regions of proteins interact with multiple binding partners. Some of the molecular switches demonstrated act as redox sensors might be of particularly interest to the reader.

### 6.1. Selenoproteins S (SeS)

Selenoprotein S (SeS), a single-pass transmembrane enzyme (21 kDa, 189 residues) containing rare amino acid selenocysteine (Sec), is implicated in many important critical processes in a cell by providing first line of antioxidant defense system by direct detoxification of reactive species, regulating sulfur-based redox pathways, inflammation, signal transduction, calcium influx, and maintain intracellular redox homeostasis via regulation of stress [30,31]. Interestingly, dislocation of improperly folded proteins from the ER to the cytoplasm for degradation also desires SeS. *In vivo*, SeS is synthesized by its own special tRNA^[sec]^. A highly conserved stem-loop SEC insertion sequence (SECIS) element at the 3’ UTRs is present on selenoprotein’s mRNAs which is necessary for UGA recognition that function as a Sec codon, not as a stop codon [32]. SECIS is modulated by two additional phylogenetically conserved stem-loop structures.

SeS contains a short segment (27 residues) resided in the ER lumen with an extended segment (132 residues) in the cytoplasm[33].NMR spectroscopy reveals that the Sec C-terminal domain (region 123-189)is intrinsically disordered in the oxidized and reduced states and enriched in charged and polar residues such as glycine, proline, and lysine plus arginine residues in 21%, 10% and 19%respectively [31,34,35]. An intra-molecular selenylsulfide bond is formed in the disordered region between Cys^174^ and Sec^188^which is reduced by the thioredoxin and thioredoxin reductases showing reduction potential of −234 mV [36]. A studyby Christensen et al. (2012) speculates that residues163–189contain substrate recognition elements which interacts with cellular redox regulator such as thioredoxin [34]. SeS is a diverse scaffolding protein, whose transmembrane (TM) helix and coiled region carry two extended α-helices which are responsible for homo/hetero-oligomerization as well as mediating protein-protein interactions [37]. The plastic nature of SeS make it suitable for fusing with 200 protein partners including oligosaccharyl transferase, multi synthetase, and nuclear pore complexes [37]. This extensive network of interactions played key roles in regulating the shape of ER, lipid metabolism as well as management of lipid droplets as any defect in the regulation would lead to cancer, inflammation autoimmune thyroid diseases, cardiovascular diseases, metabolic disorders and diabetes [38–45]. It is thought that SeS also participate in PI3K/Akt signaling pathways by modulating the expression of kinases nonetheless the mechanism is largely unknown [44].

As a part of ERAD complex, this membrane residing protein contains a p97-ATPase-valosin-containing protein (VCP) interacting motif (residues 78-88) that induce binding with p97, which in turn provide energy to dissemble and pulling of misfolded proteins to the cytoplasm by the proteasome thereby, increasing cell survival by neutralizing ER stress [42,46]. The conformation of this region remains disorder even upon binding with p97. Sec form multimeric complexes also with derlin 1 and 2 (components of putative ERAD channel), UBXN 8 along with derlin-1 controls the degradation of lipidated apolipoprotein B-100, UBXN 6 tethers p97 to the ER membrane, VCP accessory protein ubiquitin conjugation factor E4A (UBE4A),involved in multi ubiquitin chain extension, kelch containing protein 2 (KLHDC-2, HCLP-1) that takes part in the regulation of the LZIP transcription factor and selenoprotein K, an enzyme with unknown function[47–51]. Numerous studies showed the SelS-dependent protective mechanisms under glucose deprivation (fasting), treatment of cell with tunicamycin (N-glycosylation inhibitor) and thapsigargin (Ca2Þ-ATPase blocker) [52,53]. SelS up-regulates and protects the ER from stress as aggregates of misfolded proteins accumulate in the ER due to under-glycosylation and formation of unspecific disulfide bonds and Ca2Þ depletion. One study recognized that SelS is down regulated in liver, adipose tissue and skeletal muscle of type II diabetes *Psammomysobesus* animal model [45].

### 6.2. Beclin 1 (BECN1) mediates

A variety of biological processes is achieved by Beclin 1such as protection, development, apoptosis, immunity, protein sorting, trafficking, tumor suppression, and increase in lifespan. BECN1 protein consists of four domains; an IDR containing BH3 homology domain (BH3D), a flexible helical domain (FHD), a coiled-coil domain (CCD) and a β-α-repeated autophagy-specific domain (BARAD). Each domain mediates IDR endowed specific function with high specificity and reversibility thereby allowing them to control protein signaling, transcription, recruitment, and translation [5]. BECN1-IDR is enriched in disorder-promoting polar charged amino acids (51%), glycine (6%) and proline (6%). Heteronuclear single quantum coherence (HSQC, 2D-NMR) exhibited that residues (1–150) in BECN1 sequence is disordered (Figure 2) when the four cysteines of invariant CXXC and CXD/EC motifs were mutated to serines [54]. BECN1-IDR consist of numerous binding motifs including anchor regions and eukaryotic linear motifs (ELMs) that are sites of PTM to regulate conformation changes and interactions [55]. BECN1-IDR contains four anchor regions where the fourth anchor extends into the FHD. Upon binding to the partners, the molecular recognition features (MoRFs) in the first two anchor region undergoes disorder-to-helix transitions whereas the third anchor region, BH3D (residues 105-130), is disordered and folds into a helix upon binding to numerous BCL2 homologs [58,59].

**Figure 1.**
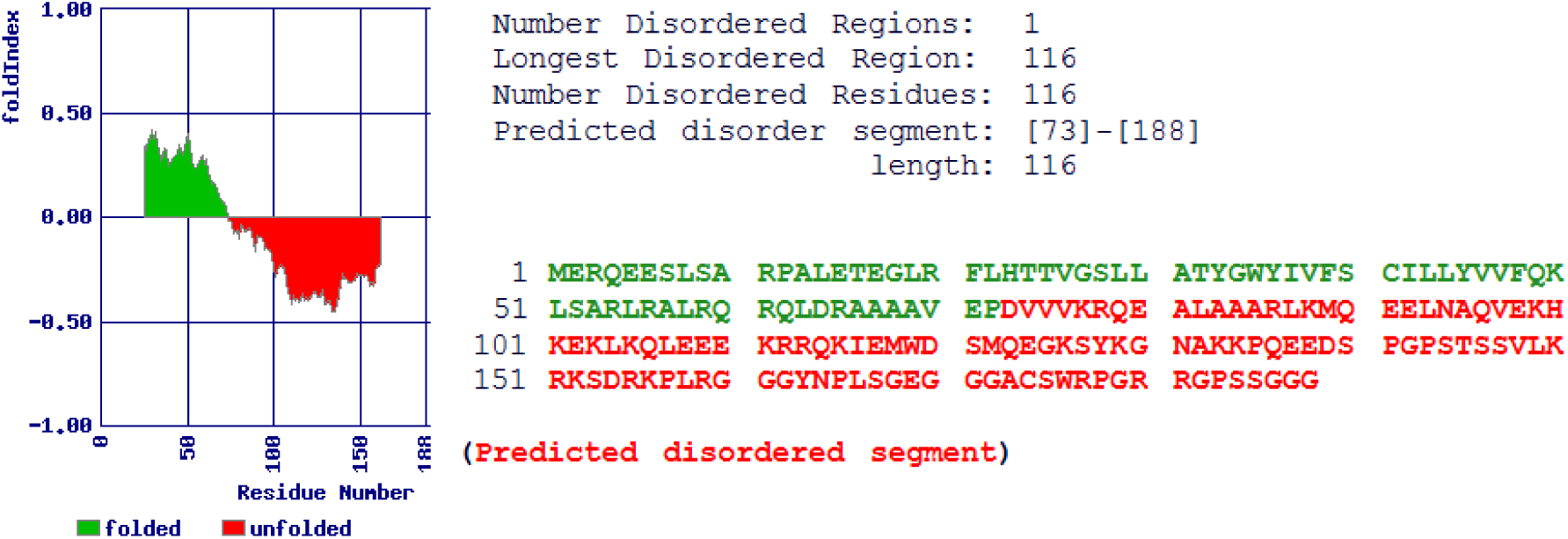
IDR detection of human SelenoproteinS (SeS) by FoldIndex software [84]

**Figure 2.**
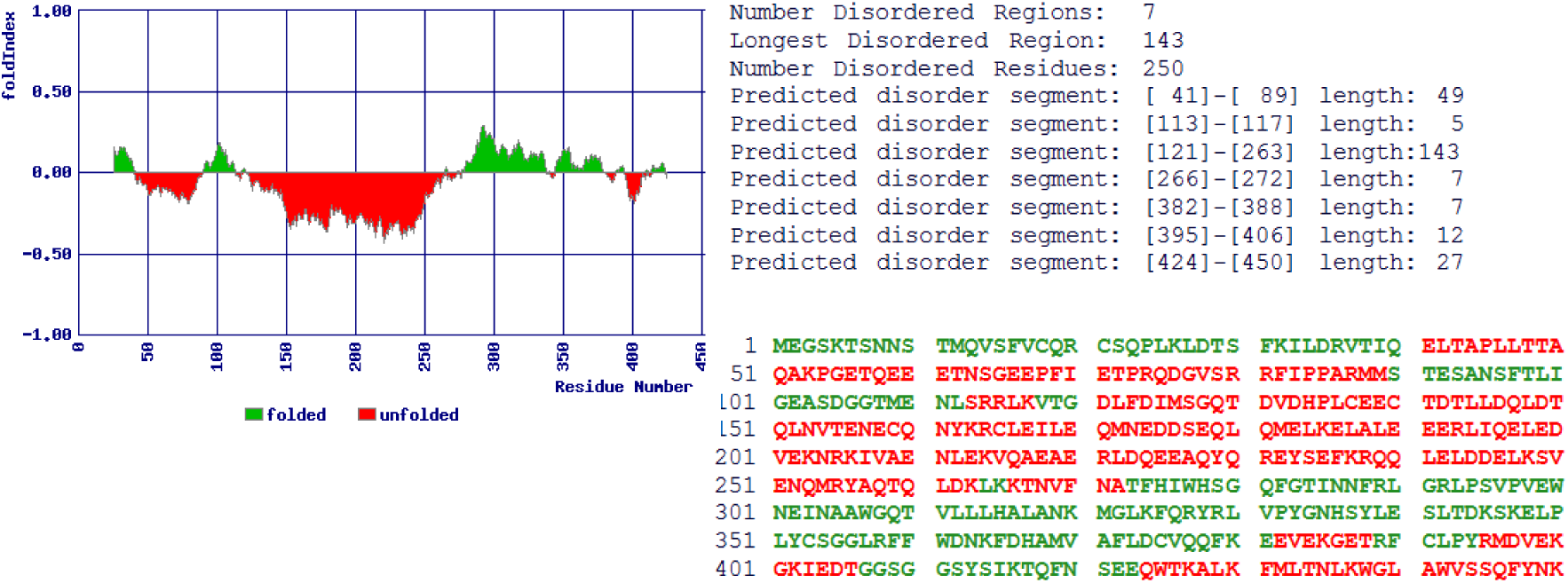
IDR detection of human Beclin-1 by FoldIndex software [84]

### 6.3. Granulins (GRNs)

Progranulin (PGRN, 68 kDa, a glycoprotein) is consisted of a signal peptide with repeats of cysteine-rich motifs. Proteolysis of PGRN by elastase and extracellular proteases yields several fragments called granulins (6-25 kDa) [60,61]. The unique small family (1-7) of GRN proteins (∼6 kDa) forming 6 intramolecular disulfide bonds, possesses 12 conserved high-density cysteines comprising (20%) the total amino acid residues. The position of cysteines in all GRNs have a single cysteines at both C- and N-termini [62]. GRNs diverse roles include wound healing, embryonic growth, inflammation and a slightly defect in their regulation would lead to neurodegenerative disorders [62]. Interestingly, granulins displayed antagonistic function e.g., GRN-4 promotes proliferation in A431epithelial cell line while GRN-3 inhibits the growth. GRN-3, governs varying disorder propensities (Figure 3), at low concentration, is monomeric (reduced from), conversely, it experiences, at high concentration, dimerization to form a “fuzzy” complex with no defined structure (a hallmark of some IDPs) [63]. In human neuroblastoma cells GRN-3 (reduced form) activate NF-κB in a concentration-dependent manner and abrogation of the disulfide bonds renders the protein disordered dictating importance of disulfide bonds for determining overall conformation of the protein and monomer–dimer dynamics [64]. The native oxidized form of GRN-3 has irregular conformation (loops) linked jointly only by disulfide bonds, lack regular secondary structure which induced noteworthy thermal and structural stability to the protein, elaborating a exclusive adaptation of IDPs [63].

**Figure 3.**
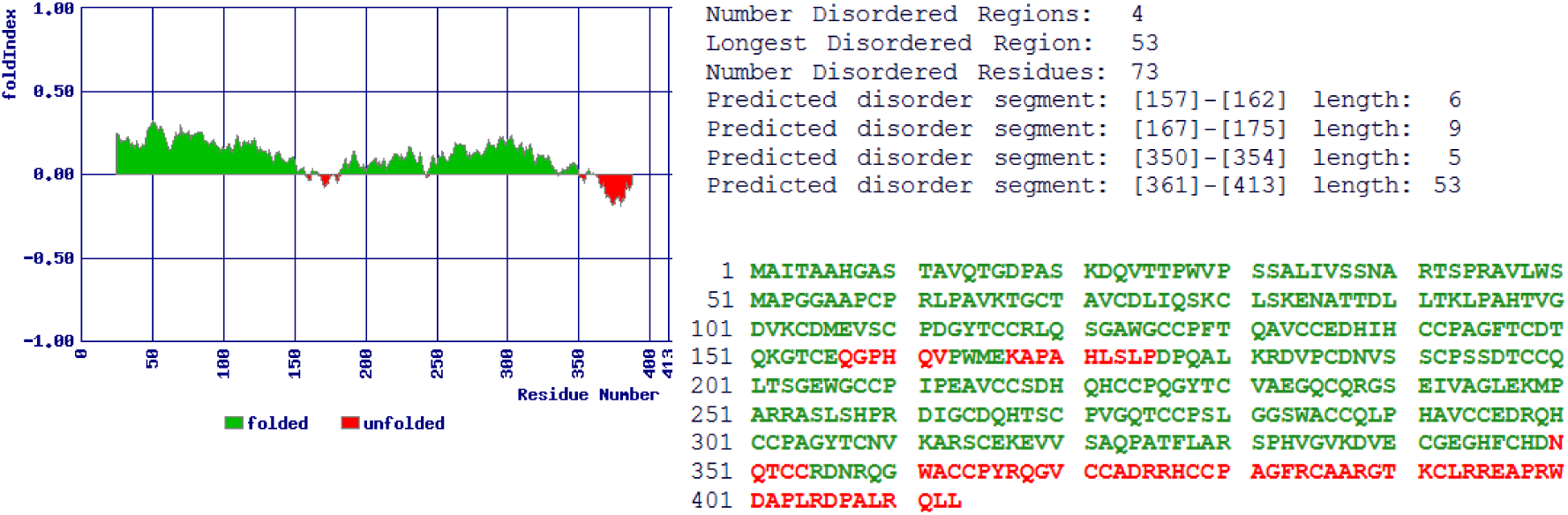
IDR recognition of human Isoform 3 of Granulins by FoldIndex software [84]

### 6.4. Cystathionine β-synthase (CBS)

CBS is included in a large family of type II pyridoxal 5’-phosphate-dependent enzymes that are regulated by gases especially with binding CO, NO and heme component. Upon changes in the redox state of a cell, the heme functions as a sensor by altering the enzymatic activity. In addition, it is involved in cellular production of gaso-transmitter H_2_S and also responsible for the sulfur metabolism [65–67]. Aberrant function of CBS has been reported for various pathological conditions such as neurodegenerative and cardiovascular disorders as well as cancer rendering it an interesting drug target [68,69]. CBS contains heme-binding motifs (HBM) or heme regulatory motifs (HRM) at the N-terminus (residues 1–70) followed by highly conserved catalytic core and a C-terminus regulatory domain. Canonical heme is bound moderately via transient interactions at C-52/H-65. However, the first 42 residues were missing in X-ray structures of CBS and this truncation reduces but does not abolish the activity of the enzyme. NMR-spectra evident that N-terminal residues (1-42) constitute an intrinsically disordered region (IDR) that interacts and contributes to the non-canonical heme scavenging site. Heme binding site in CBS at position C-15/H-22 increases CBS efficacy by approx. 30% by providing a secondary independent binding site leading to a hexa-coordinated complex which can act as trigger for other protein interactions [70,71]. This site can be an attractive drug target without losing the whole biological role [72].

### 6.5. A2B2-GAPDH

Inactivation of the Calvin cycle in land plants at night is controlled by glyceraldehyde-3-phosphate dehydrogenase (GapAB) and CP12. Gene duplication in GapA has resulted in GapB isoform that differs from GapA by a specific C-terminal extension (CTE) acquired from CP12. This CTE is responsible for thioredoxin-dependent light/dark regulation. The GapB GAPDH subunit and other proteins can be regulated by their IDR (Figure 5) [73]. Upon oxidation, a disulfide bridge is formed by two cysteines residues of the CP12-like tail, where NADP cofactor binding site is blocked due to E362 (glutamate residue) within the active site and stabilized by electrostatic interaction with R77 (arginine residue). Consequently, A2B2-GAPDH NAPDH dependent activity is inhibited due to NADPH is unable to approach the catalytic site [74]. Upon reduction, during the night-day transition, the disulfide bridge is reduced by thioredoxin f (TRX) maintaining the C-terminal extension into the active site. Thus, releasing the CP12-like tail and resuming the activity of A2B2-GAPDH [74].

**Figure 4.**
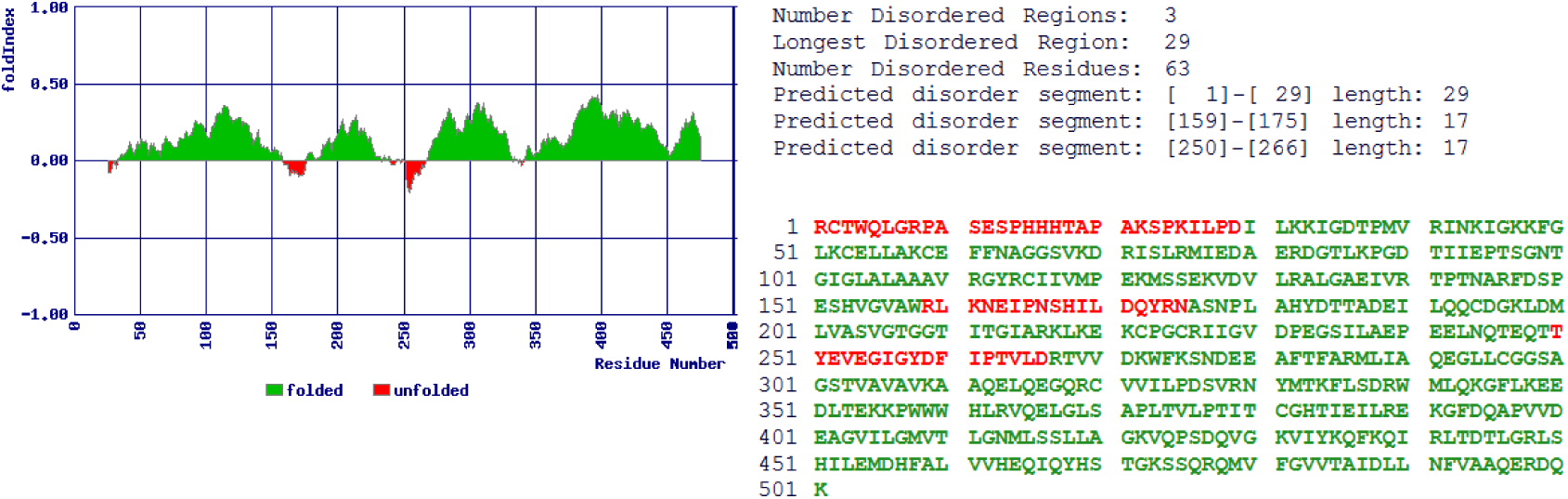
IDR detection of human Cystathionine beta-synthase (CBS) by FoldIndex software [84]

**Figure 5.**
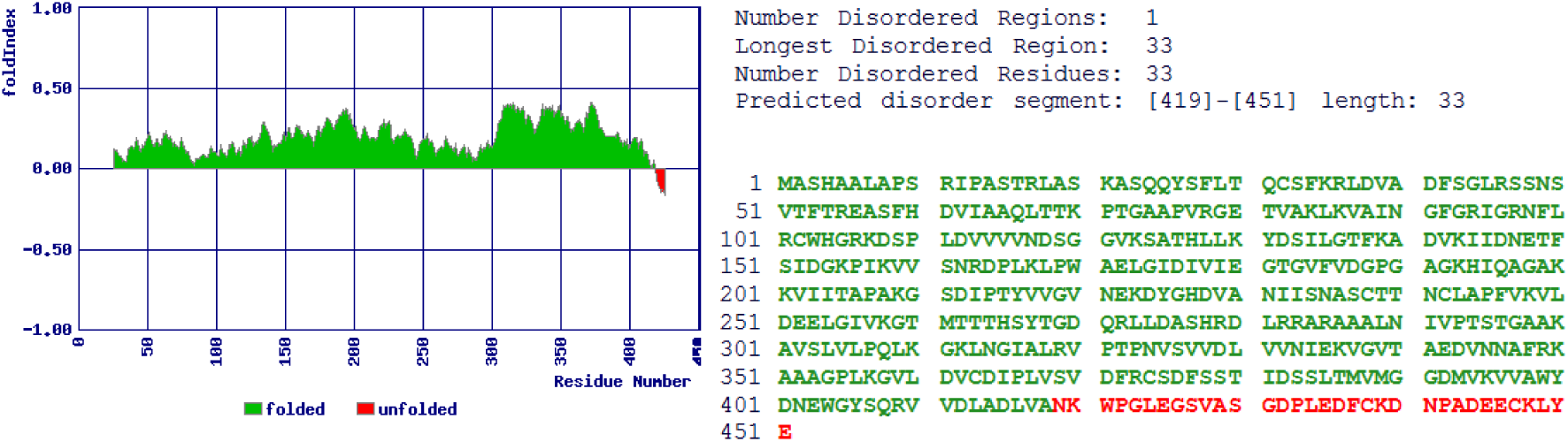
IDR of Spinacia oleracea chloroplastic Glyceraldehyde-3-phosphate dehydrogenase B form detected by FoldIndex software [84]

### 6.6. *Bdellovibrio bacteriovorus* Bd0108 protein

*Bdellovibrio bacteriovorus*, a predatory δ-proteobacterium, undergo two types of cell divisions depending on the environmental conditions; host-independent (HI, rare) and host-dependent (HD, usual). HI growth occurs on protein-rich media (in the absence of prey cells by down-regulating HD genes) while in HD state it infects and preys on gram negative (*Escherichia coli, Salmnonella, Pseudomonas auroginosa*) and gram positive pathogenic strains (*Staphlococcus aureus*) therefore, could be a potential candidate in biotechnology to control human pathogens by acting as an antimicrobial agent [75]. Remarkably, research studies reveal that the switching mechanism between HI and HD lifestyles is driven by an IDP, Bd0108, crucial for pilus regulation, predation signaling and survival [76]. Bd0108, a monomeric protein (101 residues), is needed for type IVb pilus formation and aids in the connection to gram-positive/negative bacteria for successive invasion [77,78]. Type IV pili and its associated regulatory genes (Bd0108 and Bd0109) function in conjugation, secretion of proteins at surface and cell-cell adhesion [79,80]. Mutant Bd0108 strains lack pilus, can’t invade the prey, deficient in HD stage and transformed to HI cycle [78,81,82]. It is considered that a growth mode signal is send by pilus extrusion/retraction depending on Bd0108 expression to invade inside bacteria or not indicating IDPs played crucial role in the life cycle of an organism, a noteworthy extension to IPDs functions [78].

Bd0108 sequence, N-terminal region (24-66 residues), consists of 8 out of 10 hydrophobic residues and exhibits many conserved charged and polar amino acids that can undergo multiple states. Under HD cell cycle, a potential pilus regulatory “fuzzy” complex is formed between Bd0108 (N-terminal region) and its binding partner Bd0109 that helps in predation [19]. NMR and CD studies revealed that Bd0108 exists in solution almost exclusively as a random coil and conserve intrinsically disordered nature upon binding with Bd0109 (Figure 6). It showed propensity of conformational plasticity from random coil to α-helix at its core (56–66 residues) showing oxidized thiol group and formation of a disulfide bond C-56/C-59. Four positions (CXXC motif), are responsible for separation of these residues, and their side-chains would be appeared nearby on the same surface of an α-helix, suggesting Bd0108 disulfide function as an anchor when encountering an unknown binding partner. In case of E. coli, when Bd0108 and −09 are co-expressed, they tend to localize the periplasmic regions which sense prey as well as systematize elements of entering mechanism for predation [78,83].

**Figure 6.**
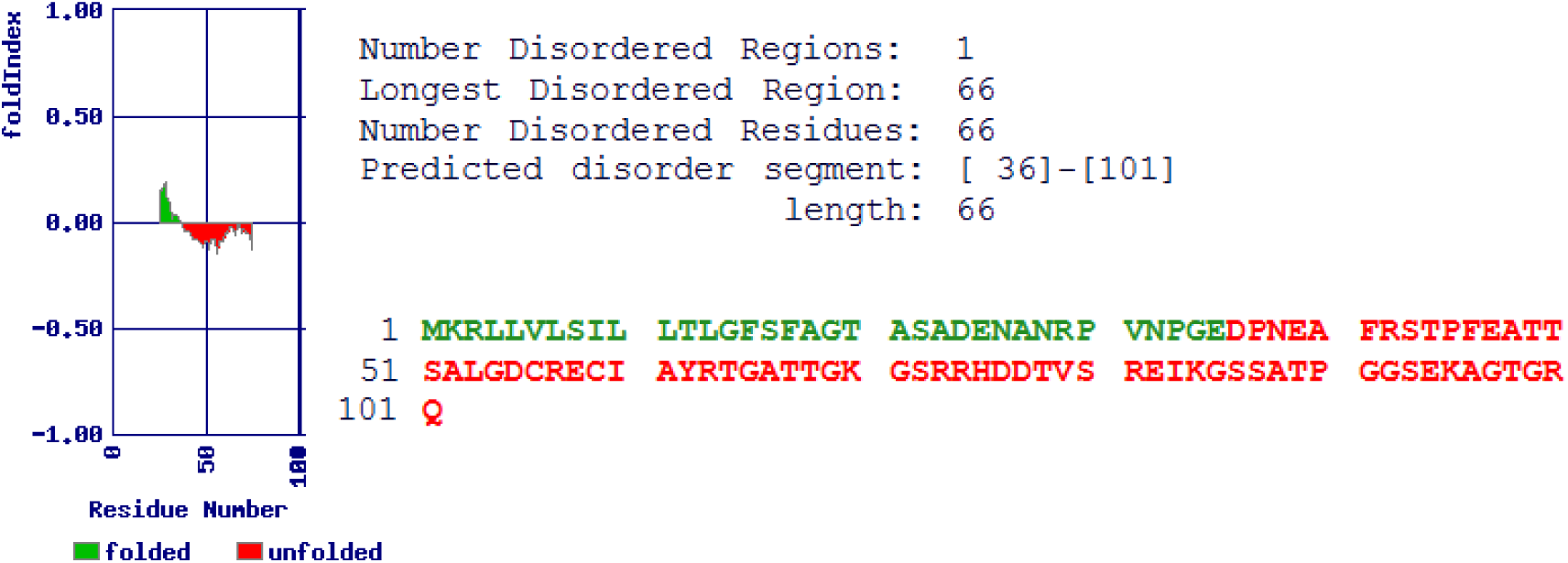
IDR of *B. bacteriovorus* (Bd0108) by FoldIndex software [84].

## Classification of redox active IDPs based on mechanism of action

i). Expulsion of transition metals (Zn, Fe, Ni, Cu) e.g., Hsp-33

ii). Reorganization of the portions of the proteins

iii). Structural and conformational changes (GAPDH/CBS/SeS)

It is also possible that in an individual IDP more than one mode of action may be employed as they behave in a different way under constant changing and challenging conditions within a cell. Owing to their structural flexibility, IDPs can be molded into specific shape depending upon the interacting partners and may undergo conformational changes that confer their diverse functions according to cell demands. It is important to whom they bind as it changes the structure and function of the protein moreover, continuous cellular motion IDRs facilitates the interaction with other proteins.

## 7. Conclusion and future perspective

Some of the functions of redox active intrinsically disordered cysteines and their mechanism of action are described in this review. The extraordinary redox status of cysteines is not only limited to provide structural and thermal stability to intrinsically disordered proteins including Grn-3 but also involved in antioxidant, anti-apoptotic and ER stress regulating activity such as in case of selenoprotein. Owing to their plastic nature they displayed different conformational and binding capacities in beclin1, help in determining lifestyle and predation in *B. bacteriovorus* and prevent pathological conditions by increasing binding capacities of CBS enzyme. Therefore, it was necessary to shed light on some of the mechanism of action of intrinsically disordered cysteines that act as molecular redox switches. Experimentally investigating the structural roles of cysteines in IDPs is not only challenging but their classification and plastic nature to form multiple complexes with unlimited number of clients make them interesting candidates for investigation and application in modern biology and medicine. Although the active roles of intrinsically disordered and their characterization is challenging due to different conformational changes, redox-sensitive regions based on different disorder tendencies helps us understand potential roles in treating many incurable diseases.

